# A machine learning approach for Colles’ fracture treatment diagnosis

**DOI:** 10.1101/2020.02.28.970574

**Authors:** Kwun Ho Ngan, Artur d’Avila Garcez, Karen M. Knapp, Andy Appelboam, Constantino Carlos Reyes-Aldasoro

## Abstract

Wrist fractures (e.g. Colles’ fracture) are the most common injuries in the upper extremity treated in Emergency Departments. Most patients are treated with a procedure called Manipulation under Anaesthesia. Surgical treatment may still be needed in complex fractures or if the wrist stability is not restored. This can lead to inefficiency in constrained medical resources and patients’ inconvenience. Previous geometric measurements in X-ray images were found to provide statistical differences between healthy controls and fractured cases as well as pre- and post-intervention images. The most discriminating measurements were associated with the texture analysis of the radial bone.

This work presents further analysis of these measurements and applying them as features to identify the best machine learning model for Colles’ fracture treatment diagnosis. Random forest was evaluated to be the best model based on validation accuracy. The non-linearity of the measurement features has attributed to the superior performance of an ensembled tree-based model. It is also interesting that the most important features (i.e. texture and swelling) required in the optimised random forest model are consistent with previous findings.

## 1 Introduction

Colles’ fracture is one of the most common fractures of the radial bone leading to posterior displacement of distal fragments at the wrist [9]. The fracture generally causes acute pain and swelling. It may even lead to residual impairment in hand and wrist motion if left untreated [14, 23].

Current evaluation of fracture severity and stability is primarily based on radiographs such as X-ray images before and, when needed, after intervention [14]. A typical radiographic study will include analysis from multiple views (i.e. posteroanterior and lateral views) of the fractured wrist. Oblique views might also be needed to better define the fracture location.

The type of treatment generally depends on fracture displacement, angulation and shortening [14]. Fracture treatment has evolved over the past two decades improving stability and anatomical functionality. The advancement has also reduced the risk of complications such as neuropathies, arthrosis, tendon ruptures and finger stiffness during the rehabilitation following a distal radius fracture [23, 10]. Minor fractures (such as extra-articular, stable and minimally displacement with no comminution) are generally treated at the Emergency Department using closed reduction and immobilization technique (e.g. manipulation under anaesthesia (MUA) [17, 3]. More complicated fractures would require subsequent surgical treatment (e.g. open reduction and internal fixation (ORIF)).

Despite considerable research outcomes [3, 2, 4, 12, 16] have been published on the appropriate treatment procedure according to fracture characteristics, the choice of treatment remains highly subjective to the X-ray interpretation by the radiologist and largely depends upon the available clinical information on a case basis.

This work aims to develop a classification model based on a dataset of X-ray images taken in the Emergency Department to detect fractured wrist cases from the healthy controls. For the fractured cases, it will further determine the success of closed reduction treatment (i.e. MUA) at pre- and post-intervention conditions.

## 2 Materials and Methods

### 2.1 Patient Dataset

A data collection of 161 independent cases of wrist fracture was used in this work. The data was sourced ethically from the Royal Devon and Exeter Hospital with Caldicott Guardian approval. Prior informed consent was obtained from each participant. The data was subsequently anonymised according to the ethics policy of the donating institution.

Each case included basic anonymised patient information (e.g. age and gender), as well as, human-annotated measurements from X-ray radiographs in posterior-anterior (PA) views. The selection of these measurements has been documented in [21]. There were 139 cases diagnosed with wrist fracture of varying severity and 22 cases of healthy control. Among the cases of fracture, it was also retrospectively classified into sub-groups of pre-successful (via MUA) (n=50), pre-unsuccessful (n=31), post-successful (n=40) and post-successful (n=18) based on patient electronic records.

### 2.2 Acquisition of X-ray Radiographs

The X-ray images were obtained with the following systems across a range of exposure factors:

1. DigitalDiagnost DidiEleva01 (Philips Medical Systems, Netherlands)
2. Mobile tablet work station (Thales, France)
3. DirectView CR 975 (Kodak, USA)
4. DirectView CD850A (Kodak, USA)
5. Definium 5000 (GE Healthcare, USA)

The raw images were stored in DICOM format [5]. Some representative cases of these images are shown in Fig. 1. These images are expected to represent the variation in image quality, positioning of wrist and presence of noisy input (e.g. collimation lines and text legends).

**Fig. 1.**
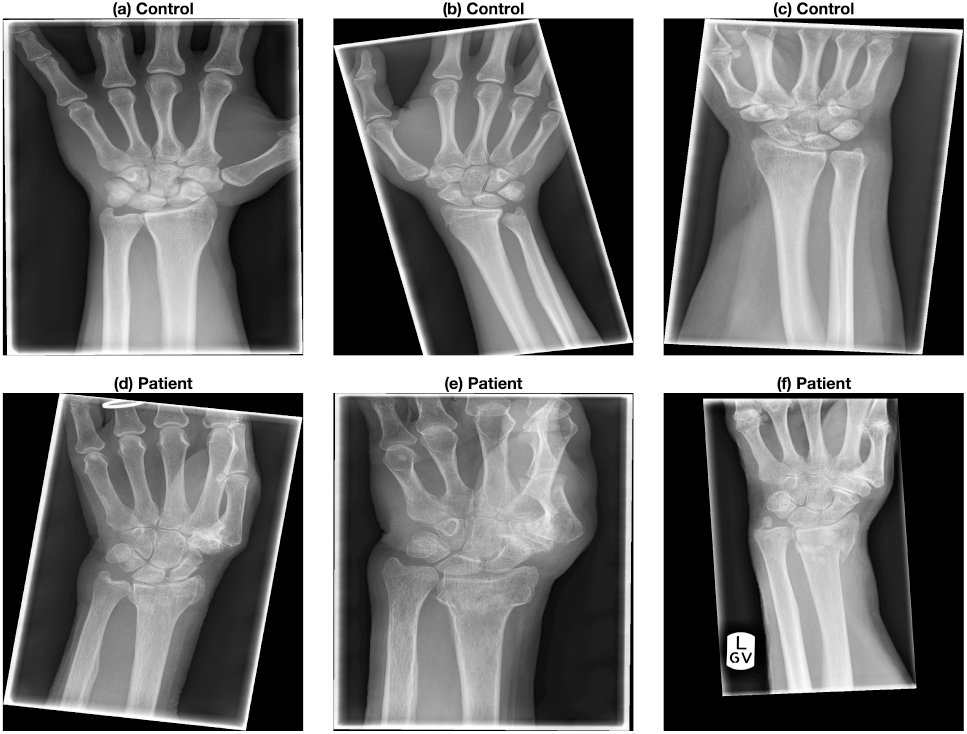
Six representative X-ray images were collected from previous clinical activity at Royal Devon and Exeter NHS Foundation Trust Emergency Department. These images present the variability in the quality, positioning of the arm and presence of noisy input (e.g. collimation lines and text legends). The images were anonymised while metadata such as age, date of acquisition, gender and clinical outcome were retained.

### 2.3 Image Pre-processing and Model Feature Generation

Three landmark points were firstly located manually - (1) base of the lunate, (2) extreme of the radial styloid and (3) centre of the metacarpal of the middle finger. The X-ray images were automatically pre-processed using Matlab based on the identified landmarks. A detailed description of the pre-processing procedure can be referred to [21]. This procedure is summarized as below.

Each image was firstly aligned vertically along the forearm. The lines of collimation were also removed. Fig. 2 shows some of the pre-processed images. Three indicator groups of measurements were extracted from the processed images to determine the appropriate treatment procedures. An illustration of geometric measurements from [21] is presented in Fig. 3.

**Fig. 2.**
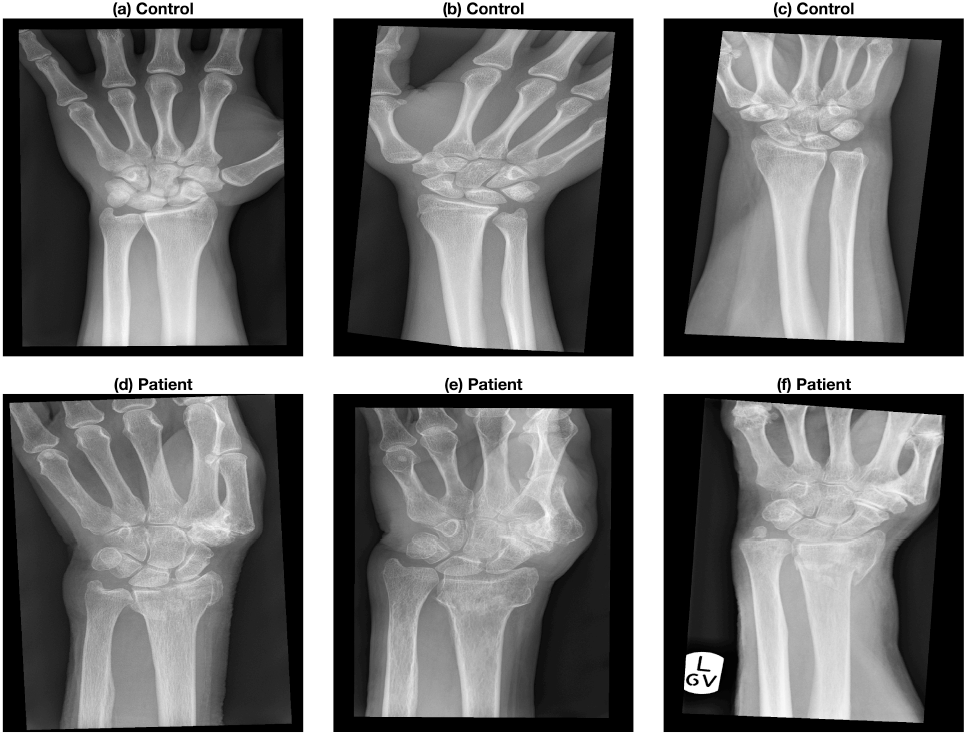
Automatic pre-processing outcome using Matlab of the X-ray images. The six representative cases shown in Fig. 1 were automatically rotated so that the forearm is vertical. In addition, the artefacts due to the collimator were removed.

**Fig. 3.**
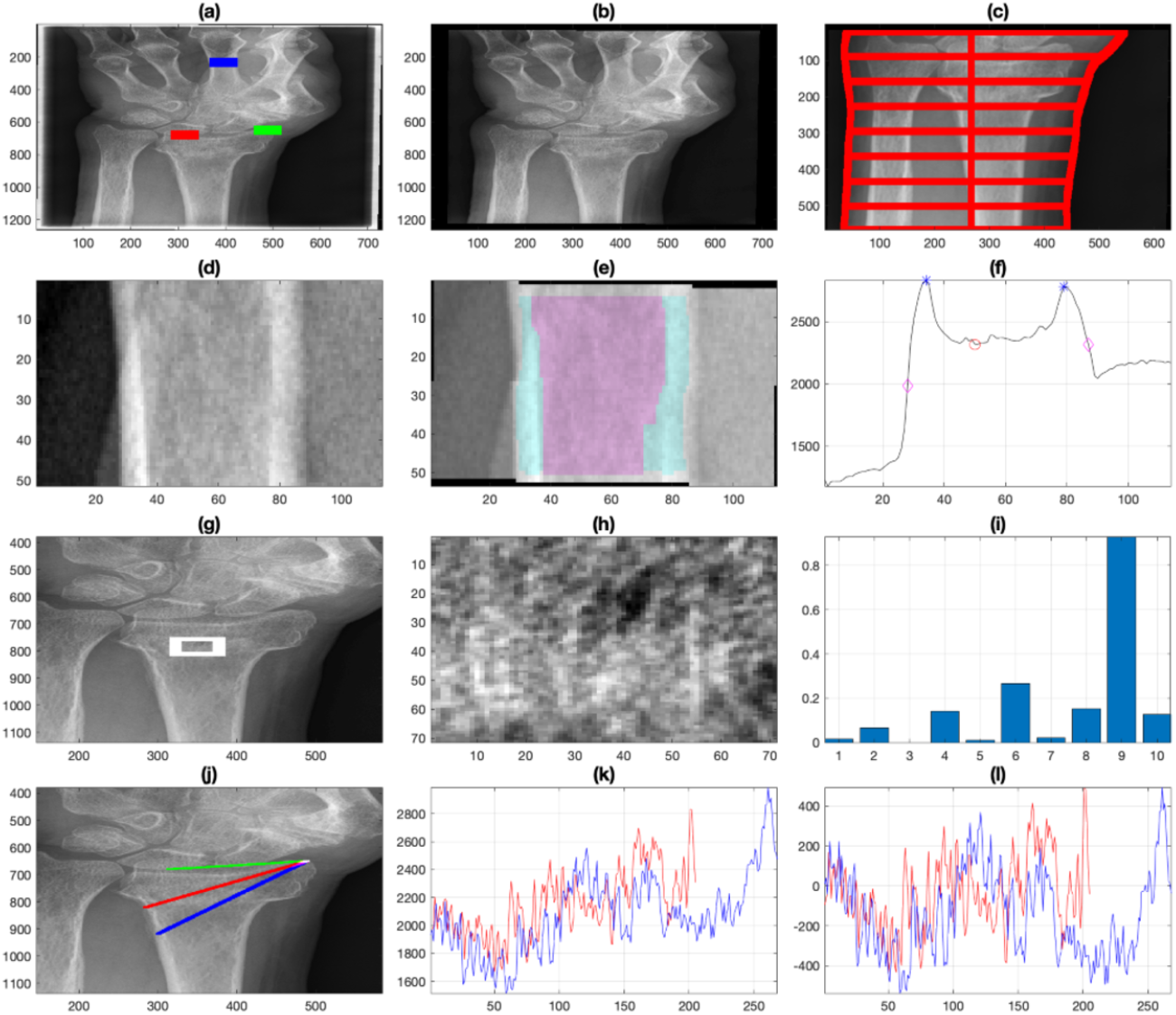
Illustration of preliminary characterisation of radiographs with geometric measurements. (a) Original radiograph with landmarks. (b) Preprocessing. (c) Boundaries and traces of forearm. (d,e,f) Cortical bone segmentation. (g,h,i) Texture analysis. (j,k,l) Intensity profile analysis. [21])

1. **Indication of Swelling:** Widths along the forearm as an indication of swelling.
2. **Indication of Osteoporosis:** Width of finger and ratio of trabecular area to the total area of interest measured at the middle finger [7, 24, 15].
3. **Texture Indication at Radial Bone:** Texture measurement from a selected region of radial bone using Local Binary Pattern (LBPs) [19]. Grayscale intensity profile across landmarks #2 and #3 to indicate the changes in texture in the radial bone.

This forms a feature set of 36 variables for model construction including age, gender and 34 image-based measurements in the three indicator groups. Table 1 presents the exact features used in this study.

**Table 1:** Grouping of Features for Treatment classification by Indicator Type.

| Grouping of Features for Treatment Classification |  |  |  |
| --- | --- | --- | --- |
| Basic Patient Info | Indication of Swelling | Indication of Osteoporosis | Texture Indication at Radial Bone |
| Age | Ratio: Width Line 1 / Width Line 4 (WW1) | Ratio: Trabecular Area / Total Area (TrabecularToTotal) | Local Binary Pattern 1 (LBP1) |
| Gender | Ratio: Width Line 2 / Width Line 4 (WW2) | Finger Width (WidthFinger) | Local Binary Pattern 2 (LBP2) |
|  | Ratio: Width Line 3 / Width Line 4 (WW3) |  | Local Binary Pattern 3 (LBP3) |
|  | Ratio: Width Line 5 / Width Line 4 (WW5) |  | Local Binary Pattern 4 (LBP4) |
|  | Ratio: Width Line 6 / Width Line 4 (WW6) |  | Local Binary Pattern 5 (LBP5) |
|  | Ratio: Width Line 7 / Width Line 4 (WW7) |  | Local Binary Pattern 6 (LBP6) |
|  | Ratio: Width Line 8 / Width Line 4 (WW8) |  | Local Binary Pattern 7 (LBP7) |
|  | Ratio: Min Width Line / Max Width Line (MinMaxWidthAtCM) |  | Local Binary Pattern 8 (LBP8) |
|  | Ratio: Width Line 1 + 8 / Width Lines 4 + 5 (W1W8/W4W5) |  | Local Binary Pattern 9 (LBP9) |
|  | Ratio: Width Line 1 + 2 / Width Lines 7 + 8 (W1W2/W7W8) |  | Local Binary Pattern 10 (LBP10) |
|  |  |  | Average Row LBP (stats.row_LBP) |
|  |  |  | Average Column LBP (stats.col_LBP) |
|  |  |  | Slope First Profile Full Line (stats.slope_1) |
|  |  |  | Slope Second Profile Full Line (stats.slope_2) |
|  |  |  | Slope First Profile Short Segment (stats.slope_short_1) |
|  |  |  | Slope Second Profile Short Segment (stats.slope_short_2) |
|  |  |  | Standard Deviation First Profile (stats.std_1) |
|  |  |  | Standard Deviation Second Profile (stats.std_2) |
|  |  |  | Standard Deviation First Profile Adjusted (stats.std_ad_1) |
|  |  |  | Standard Deviation Second Profile Adjusted (stats.std_ad_2) |
|  |  |  | Distance First Profile (dist1) |
|  |  |  | Distance Second Profile (dist2) |

### 2.4 Predictive Classification Models

Three different classification models were used for the selection of the best predictive model - Logistic Regression (LR), Decision Tree (DT) and Random Forest (RF). These models have been widely used in classification for medical use cases [1, 6, 8, 11] and have found varying success in prediction.

Logistic Regression is a linear model for classification based on the probabilities describing the possibility of outcomes (i.e. classes) of a single sample using a logistic function [20]. In a multi-class scenario, the class with the highest probability denotes the predicted class [22]. Decision Trees are non-parametric classification models by learning simple decision rules from features to predict class value [20]. Random Forest is an ensemble classifier by fitting a pre-defined number of decision trees and finding the most probable class from the average of the leaf nodes of the decision trees [20, 18].

Given that the number of samples (n=161) in the dataset is limited, upsampling has been applied in this study by data replication using sampling with replacement (random state: 0). The resulting sample dataset has a size of 10000 samples. The distribution of the 5 classes in the upscaled dataset remains consistent. This is based on the assumption that the dataset of X-ray images is representative of the distribution of procedure prescribed within a typical Emergency Department and has a natural bias towards fractured cases.

This work was written in Python and used the Scikit-Learn package for model construction and evaluation. Each model was trained and evaluated in a 10-fold cross-validation. Default values for model hyperparameters can be assumed unless otherwise stated in the model evaluation. The average accuracy of each model was used for performance comparison. All models were set to a fixed random state (i.e. 1 for this study) to ensure result reproducibility.

## 3 Analysis and Evaluation

### 3.1 Exploratory Data Analysis

The cross-correlation between the given features was first investigated. A heatmap (Fig. 4) was generated to determine the degree of correlation. A cell of red between two features indicates a positive correlation while a cell of blue indicates a negative correlation. The colour intensity represents the strength of the correlation. It can be observed that there is a strong correlation between features within indicator groups (e.g. wrist width ratios as swelling indicators, local binary profiles and grayscale intensity profiles on the texture of the radial bones).

**Fig. 4.**
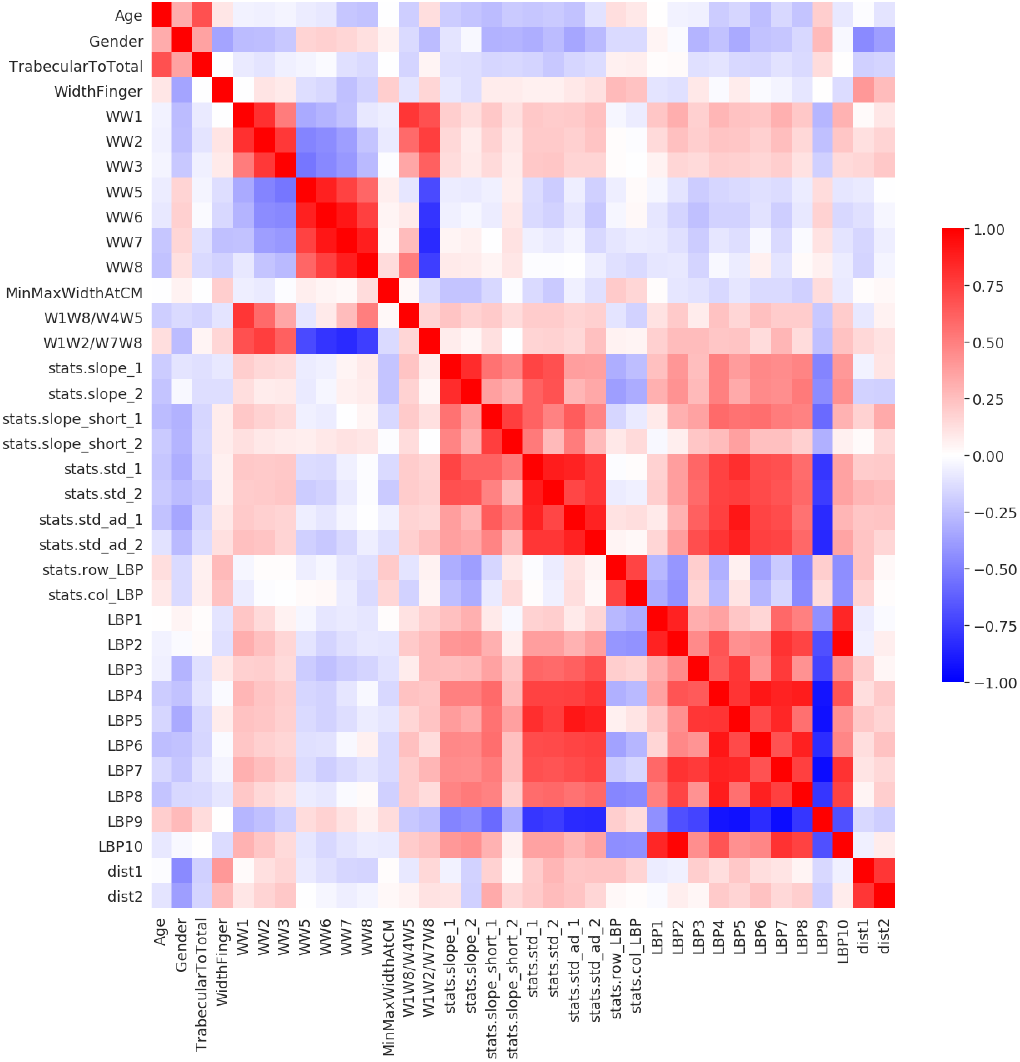
Heatmap showing clusters of cross-correlations among variables in the Fracture dataset. Blue indicates a negative correlation and red indicates a positive correlation. The colour intensity of each cell shows the strength of the correlation.

Interestingly, LBP9 has a distinctive negative correlation with other texture related features. Given the bin value range for LBP9 (approx. 204 to 229 in binary format), it may indicate the contrasting presence of edges (e.g. hairline cracks) and smooth surface (e.g. blocks of whiteness of a cast).

In addition, the low cross-correlation across different indicator groups suggests that these groups were relatively independent. This was useful to assess the significance of the indicator groups on classification performance that will be discussed later in this work.

To understand the univariate correlation for the classification task, distribution plots of all the features (except LBP3 & LBP5) are presented in Fig. 5. LBP3 and LBP5 were omitted due to the predominant values of zeros (0) across the cases causing a distribution calculation error. It can be observed that some features (e.g. Width of Finger) showed no significant difference across classes while others (e.g. Local Binary Pattern) showed different distribution between pre-treatment and post-treatment conditions. Age and Radial Bone Landmark Distance (dist 2) also showed differences between healthy wrist and fractured wrist. No feature showed a significant difference in distribution between successful and unsuccessful treatments. These findings were consistent with the findings published in [21].

**Fig. 5.**
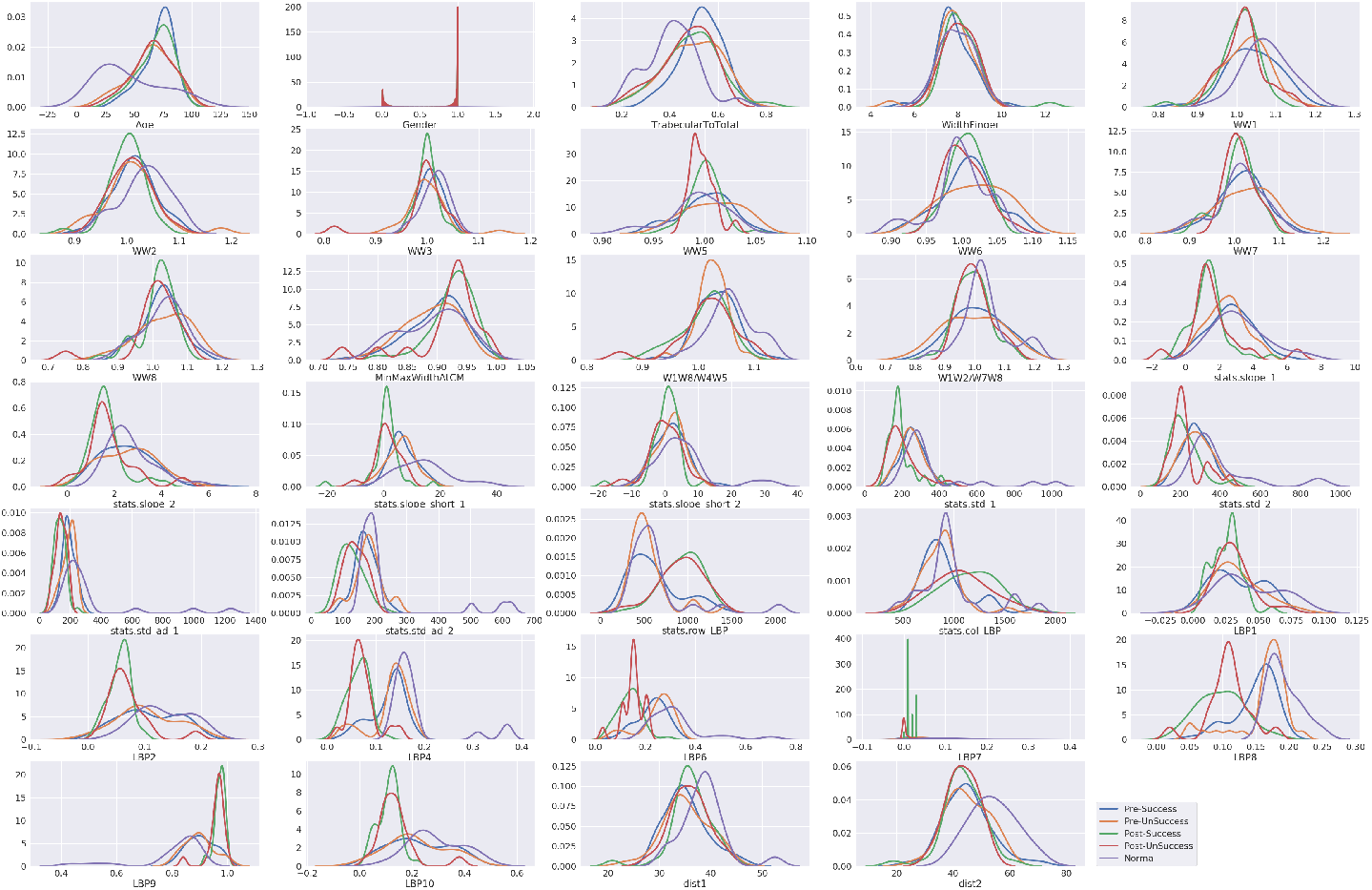
Univariate distribution by treatment classes to identify prominent features for classification.

### 3.2 Model Evaluation

Based on the initial exploratory data analysis, univariate correlation for classification had been poor. This had implication to the model performance accuracy used in this work. Fig. 6 shows the summary results of the 3 models over 10-fold cross-validation. The poor univariate correlations and linear separability could be attributing to the poor performance for the logistic regression model (Ave. Acc.: 71.0% & weighted F1-Score: 0.69).

**Fig. 6.**
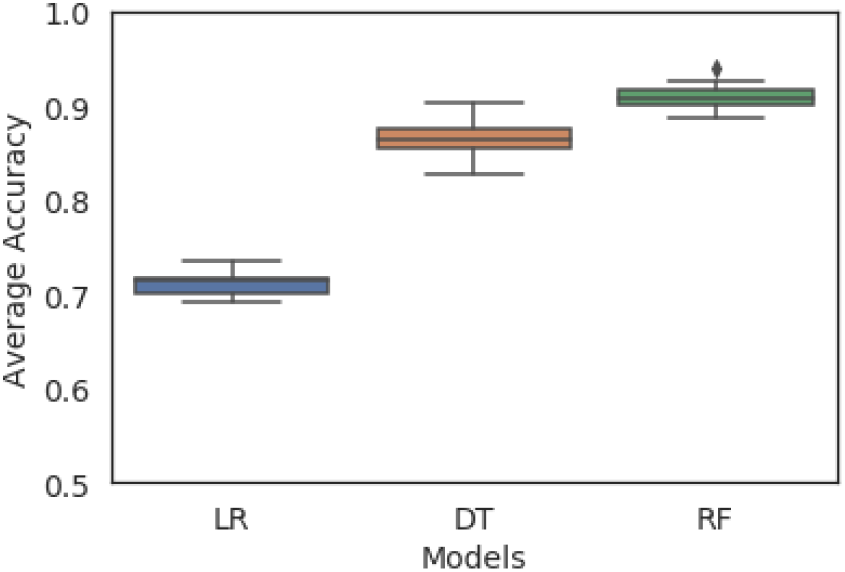
Classification Performance for Logistic Regression (LR), Decision Tree (DT) and Random Forest (RF) models over 10-fold cross-validation for prediction accuracy.

The accuracy of a decision tree showed significant improvement (Ave. Acc.: 86.7% & weighted F1 Score: 0.88). The tree branches allowed better classification through non-linear separation. A maximum tree depth of 6 (i.e. square root of the total number of features) was used according to [13]. Fig. 7 presents a sample distribution plot of a leaf node in a decision tree where post-treatment samples were completely separated and a threshold can be seen for separating the healthy samples from the fractured pre-treatment samples.

**Fig. 7.**
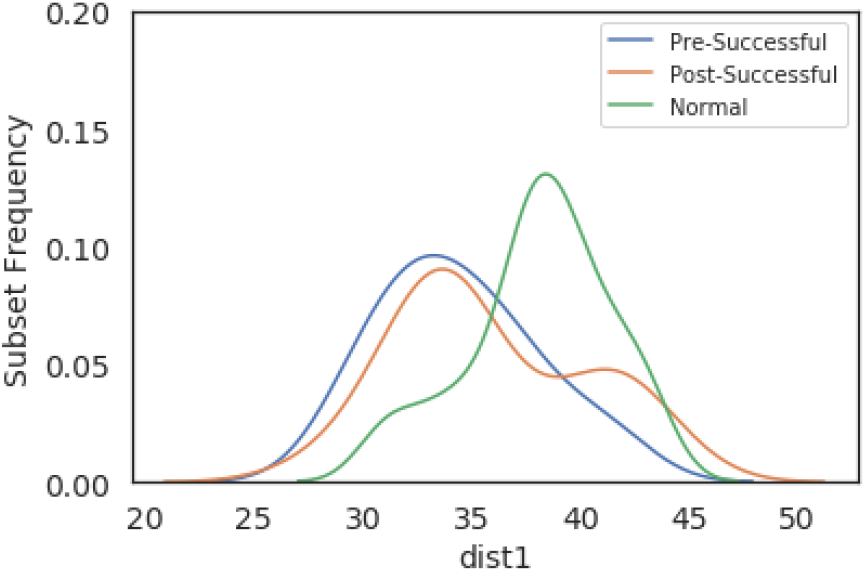
Distribution of radial bone landmark distance (dist_1) within a leaf node (Local Binary Pattern 4 > 0.09 & Local Binary Pattern 8 > 0.16 & Ratio of Wrist Line 2 over Wrist Line 4 <= 1.04).

In order to further boost the accuracy of the model, a random forest of 4 decision trees was used with the same tree hyperparameters as the decision tree model. The ensemble classifier has demonstrated an improvement of more than 4% to an accuracy of 91.0% (weighted F1 Score: 0.87). The aggregation of weak classifications from each tree enhanced the probability of the most expected class. The minor decline in F1 score was primarily attributed to the poor discrimination of procedure success in post-intervention condition with a small input sample size.

The impact of different indicator groups on classification accuracy was also investigated. Fig. 8 presents the accuracy of the same configured random forest model trained using different subsets of features. It can be clearly observed that swelling indicators (RF_Swelling) and local binary patterns alone (RF_LBP_Only) predicted poorly against the base model with 36 features (i.e. RF All) due to missing information. When all texture features at the radial bone were used (RF_Texture), it achieved similar accuracy as the base model. Despite the texture features captured most of the information needed for the classification task, this resulted in a large variation in accuracy during cross-validation subjecting to the random choice of features selected for each tree in the random forest model.

**Fig. 8.**
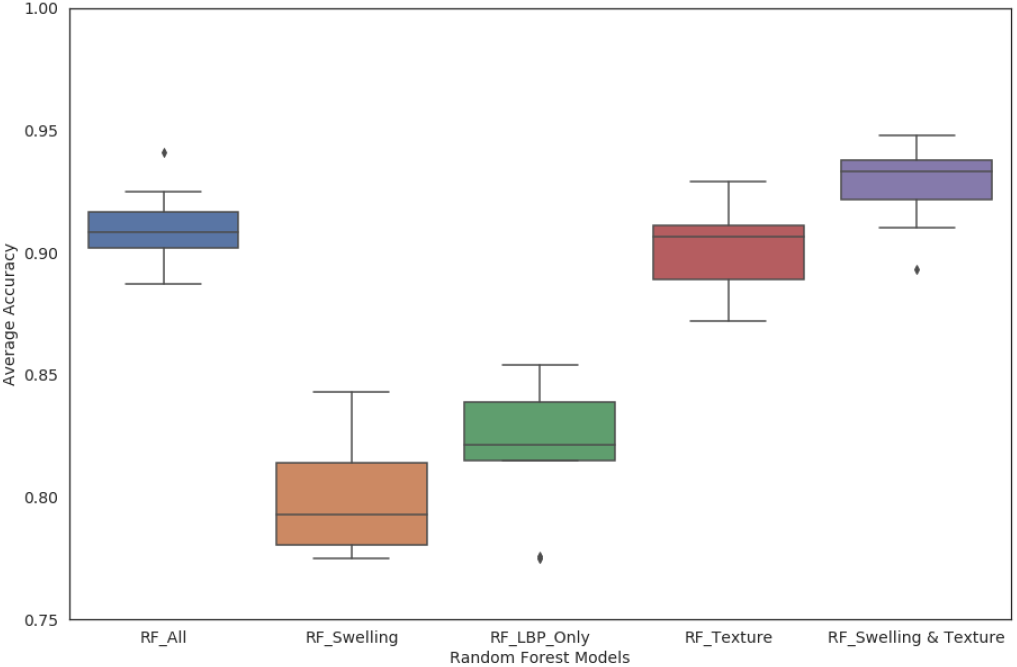
Comparison of Random Forest Models based on different category groups. (a) All Features (RF All), (b) All Swelling Features (RF Swelling), (c) Local Binary Patterns Only (RF LBP Only), (d) All Texture Features (RF Texture), (e) All Swelling & Texture Features (RF Swelling & Texture).

In the final model (RF_Swelling & Texture), other indicator groups (e.g. swelling) were used to provide supplemental information with an aim to boost accuracy. The better performance against the base model indicated that relevant supplementary features could enhance performance despite its weak predictability on its own. It however also implied that the patient’s age, gender and indication of osteoporosis were not the attributing features for this classification task and, in fact, hampered the random selection of the most important features used by the random forest model (i.e. lower accuracy observed in the base model.

## 4 Conclusion

This work has presented the complexity and multivariate nonlinearity for treatment procedure classification. Random Forest model was shown as the model of choice for this nonlinear tasks where weak classifications from decision trees were ensembled with an average prediction accuracy of 91.0%.

Based on the analysis of the input data and model outcome, this work has found similar feature importance as [21] where texture indicators at the radial bones were crucial to the classification of the input images. Local Binary Profile, together with the intensity changes across landmark points, essentially provided information on the presence of edges (i.e. signs of fracture). Other supplement information, e.g. swelling, would aid prediction but did not classify well on its own.

From these findings, it is expected that a convolution neural network (CNN) models for classification will yield similar, if not better, performance given the mechanism of feature extraction across a CNN architecture. This will form the basis for investigation in future work.

## References

1. Scikit-Learn Official Webpage, https://scikit-learn.org/stable/

2. Arora, R., Gabl, M., Gschwentner, M., Deml, C., Krappinger, D., Lutz, M.: A Comparative Study of Clinical and Radiologic Outcomes of Unstable Colles Type Distal Radius Fractures in Patients Older Than 70 Years: Nonoperative Treatment Versus Volar Locking Plating. Journal of Orthopaedic Trauma 23(4), 237–242 (Apr 2009). https://doi.org/10.1097/BOT.0b013e31819b24e9

3. Barai, A., Lambie, B., Cosgrave, C., Baxter, J.: Management of distal radius fractures in the emergency department: A long-term functional outcome measure study with the Disabilities of Arm, Shoulder and Hand (DASH) scores. Emergency Medicine Australasia 30(4), 530–537 (2018). https://doi.org/10.1111/1742-6723.12946, https://onlinelibrary.wiley.com/doi/abs/10.1111/1742-6723.12946

4. Bartl, C., Stengel, D., Bruckner, T., Rossion, I., Luntz, S., Seiler, C., Gebhard, F.: Open reduction and internal fixation versus casting for highly comminuted and intra-articular fractures of the distal radius (ORCHID): protocol for a randomized clinical multi-center trial. Trials 12(1), 84 (Mar 2011). https://doi.org/10.1186/1745-6215-12-84, https://doi.org/10.1186/1745-6215-12-84

5. Bidgood, W.D., Horii, S.C.: Introduction to the ACR-NEMA DICOM standard. Radiographics: A Review Publication of the Radiological Society of North America, Inc 12(2), 345–355 (Mar 1992). https://doi.org/10.1148/radiographics.12.2.1561424

6. Bishop, C.: Pattern Recognition and Machine Learning. Information Science and Statistics, Springer-Verlag, New York (2006), https://www.springer.com/gp/book/9780387310732

7. Bloom, R.A., Laws, J.W.: Humeral cortical thickness as an index of osteoporosis in women. The British Journal of Radiology 43(512), 522–527 (Aug 1970). https://doi.org/10.1259/0007-1285-43-512-522, https://www.birpublications.org/doi/10.1259/0007-1285-43-512-522

8. Breiman, L.: Random Forests. Machine Learning 45(1), 5–32 (Oct 2001). https://doi.org/10.1023/A:1010933404324, https://doi.org/10.1023/A:1010933404324

9. Colles, A.: On the fracture of the carpal extremity of the radius. The New England Journal of Medicine, Surgery and Collateral Branches of Science 3(4), 368–372 (1814). https://doi.org/10.1056/NEJM181410010030410, https://doi.org/10.1056/NEJM181410010030410

10. Cooney, W.P., Dobyns, J.H., Linscheid, R.L.: Complications of Colles’ fractures. The Journal of Bone and Joint Surgery. American Volume 62(4), 613–619 (1980)

11. Forman, G., Cohen, I.: Learning from Little: Comparison of Classifiers Given Little Training. In: Boulicaut, J.F., Esposito, F., Giannotti, F., Pedreschi, D. (eds.) Knowledge Discovery in Databases: PKDD 2004. pp. 161–172. Lecture Notes in Computer Science, Springer, Berlin, Heidelberg (2004)

12. Grewal, R., MacDermid, J.C., King, G.J.W., Faber, K.J.: Open Reduction Internal Fixation Versus Percutaneous Pinning With External Fixation of Distal Radius Fractures: A Prospective, Randomized Clinical Trial. Journal of Hand Surgery 36(12), 1899–1906 (Dec 2011). https://doi.org/10.1016/j.jhsa.2011.09.015, https://www.jhandsurg.org/article/S0363-5023(11)01192-0/abstract

13. Hastie, T., Tibshirani, R., Friedman, J.: The Elements of Statistical Learning: Data Mining, Inference, and Prediction, Second Edition. Springer Series in Statistics, Springer-Verlag, New York, 2 edn. (2009). https://doi.org/10.1007/978-0-387-84858-7, https://www.springer.com/gp/book/9780387848570

14. Hsu, H., Fahrenkopf, M.P., Nallamothu, S.V.: Wrist Fracture. In: StatPearls. StatPearls Publishing, Treasure Island (FL) (2020), http://www.ncbi.nlm.nih.gov/books/NBK499972/

15. Jantzen, C., Cieslak, L.K., Barzanji, A.F., Johansen, P.B., Rasmussen, S.W., Schmidt, T.A.: Colles’ fractures and osteoporosis–A new role for the Emergency Department. Injury 47(4), 930–933 (Apr 2016). https://doi.org/10.1016/j.injury.2015.11.029

16. Kapoor, H., Agarwal, A., Dhaon, B.K.: Displaced intra-articular fractures of distal radius: a comparative evaluation of results following closed reduction, external fixation and open reduction with internal fixation. Injury 31(2), 75–79 (Mar 2000). https://doi.org/10.1016/S0020-1383(99)00207-7, https://www.injuryjournal.com/article/S0020-1383(99)00207-7/abstract

17. Laseter, G.F., Carter, P.R.: Management of Distal Radius Fractures. Journal of Hand Therapy 9(2), 114–128 (Apr 1996). https://doi.org/10.1016/S0894-1130(96)80070-6, http://www.sciencedirect.com/science/article/pii/S0894113096800706

18. Marshall, R.J.: The use of classification and regression trees in clinical epidemiology. Journal of Clinical Epidemiology 54(6), 603–609 (Jun 2001). https://doi.org/10.1016/S0895-4356(00)00344-9, http://www.sciencedirect.com/science/article/pii/S0895435600003449

19. Ojala, T., Pietikäinen, M., Harwood, D.: A comparative study of texture measures with classification based on featured distributions. Pattern Recognition 29(1), 51–59 (Jan 1996). https://doi.org/10.1016/0031-3203(95)00067-4, http://www.sciencedirect.com/science/article/pii/0031320395000674

20. Podgorelec, V., Kokol, P., Stiglic, B., Rozman, I.: Decision Trees: An Overview and Their Use in Medicine. Journal of Medical Systems 26(5), 445–463 (Oct 2002). https://doi.org/10.1023/A:1016409317640, https://doi.org/10.1023/A:1016409317640

21. Reyes-Aldasoro, C.C., Ngan, K.H., Ananda, A., Garcez, A.d., Appelboam, A., Knapp, K.M.: Geometric Semi-automatic Analysis of Colles’ Fractures. medRxiv p. 2020.02.18.20024562 (Feb 2020). https://doi.org/10.1101/2020.02.18.20024562, https://www.medrxiv.org/content/10.1101/2020.02.18.20024562v1

22. Shaikhina, T., Lowe, D., Daga, S., Briggs, D., Higgins, R., Khovanova, N.: Decision tree and random forest models for outcome prediction in antibody incompatible kidney transplantation. Biomedical Signal Processing and Control 52, 456–462 (Jul 2019). https://doi.org/10.1016/j.bspc.2017.01.012, http://www.sciencedirect.com/science/article/pii/S1746809417300204

23. Simic, P.M., Weiland, A.J.: Fractures of the Distal Aspect of the Radius: Changes in Treatment Over the Past Two Decades. JBJS 85(3), 552–564 (Mar 2003)

24. Webber, T., Patel, S.P., Pensak, M., Fajolu, O., Rozental, T.D., Wolf, J.M.: Correlation Between Distal Radial Cortical Thickness and Bone Mineral Density. The Journal of Hand Surgery 40(3), 493–499 (Mar 2015). https://doi.org/10.1016/j.jhsa.2014.12.015, http://www.sciencedirect.com/science/article/pii/S0363502314017201

